# New β-propellers are continuously amplified from single blades in all major lineages of the β-propeller superfamily

**DOI:** 10.1101/2022.03.13.484165

**Authors:** Joana Pereira, Andrei N. Lupas

**Author notes:** Biozentrum and Swiss Institute of Bioinformatics, University of Basel, Spitalstrasse 41, 4056 Basel, Switzerland (Joana Pereira).

## Abstract

β-Propellers are toroidal folds, in which consecutive supersecondary structure units of four anti-parallel β-strands – called blades – are arranged radially around a central axis. Uniquely among toroidal folds, blades span the full range of sequence symmetry, from near identity to complete divergence, indicating an ongoing process of amplification and differentiation. We have proposed that the major lineages of β-propellers arose through this mechanism and that therefore their last common ancestor was a single blade, not a fully formed β-propeller. Here we show that this process of amplification and differentiation is also widespread within individual lineages, yielding β-propellers with blades of more than 60% median pairwise sequence identity in most major β-propeller families. In some cases, the blades are nearly identical, indicating a very recent amplification event, but even in cases where such recently amplified β-propellers have more than 80% overall sequence identity to each other, comparison of their DNA sequence shows that the amplification occurred independently.

## Introduction

Duplication is one of the most common mechanisms for the emergence of biomolecular novelties (Levasseur and Pontarotti, 2011). It can occur at different levels - from a small number of nucleotides up to whole genomes. When it happens at the scale of short nucleotide segments, it may lead to the amplification of subdomain-sized fragments and to the emergence of repetitive protein domains (Andrade, Perez-Iratxeta and Ponting, 2001; Söding and Lupas, 2003). Usually, the monotonous amplification of such fragments produces open-ended, solenoid folds, as in tetratricopeptide (TPR)-repeat (Blatch and Lässle, 1999) and leucine-rich repeat (LRR) proteins (Matsushima *et al*., 2005), but in some cases it may lead to closed, globular folds. One example is the β-propeller fold (Jawad and Paoli, 2002; Chaudhuri, Söding and Lupas, 2008; Kopec and Lupas, 2013; Smock *et al*., 2016) adopted by a vast family of scaffolds involved in macromolecular interactions and catalysis. β-Propeller domains are found in all kingdoms of life (Pereira and Lupas, 2021) and are involved in a wide range of biological processes (Fülöp and Jones, 1999; Pons *et al*., 2003; Guruprasad and Dhamayanthi, 2004; Chen, Chan and Wang, 2011). Their repetitive unit, the ‘blade’, is an ancestral 4-stranded antiparallel β-meander of typically 40-50 residues (Alva, Söding and Lupas, 2015), which is radially arranged around a channel to form a toroid fold, the ‘propeller’ (Andrade, Perez-Iratxeta and Ponting, 2001; Söding and Lupas, 2003). Currently, more than 300 structures have been experimentally obtained for a variety of β-propellers with 4 up to 12 blades.

Despite the high structural similarity of their blades, β-propellers span a wide range of internal sequence symmetry, from near identity (>90% sequence identity) to almost full differentiation (<20% sequence identity) (Pereira and Lupas, 2021), a unique feature and a crucial factor for their wide functional diversification. The most symmetric β-propeller known to date is that of the WRAP domain in Npun_R6612 from the cyanobacterium *Nostoc punctiforme* PCC73102, with only 10 point mutations across 14 predicted blades (Chaudhuri, Söding and Lupas, 2008; Dunin-Horkawicz, Kopec and Lupas, 2014). The near identity of its repetitive units and the absence of synonymous mutations in their encoding DNA suggests that this β-propeller was amplified very recently. The 14 blades fold into two separate 7-bladed β-propellers, but are able to produce a multitude of fold topologies when amplified to different copy numbers in the same chain (Afanasieva *et al*., 2019).

WRAP forms the C-terminal part of a protein with a STAND ATPase domain, whose gene is transcriptionally coupled to another STAND ATPase carrying an N-terminal Toll/Interleukin receptor (TIR) domain (Dunin-Horkawicz, Kopec and Lupas, 2014). This operon-like structure is conserved in other cyanobacteria, where the TIR-containing protein may also carry an additional N-terminal caspase domain and thus combines three core domains of eukaryotic innate immunity and apoptosis. The further discovery of similar proteins with recently amplified or disrupted (by frameshifts or in-frame stop codons) β-propellers, suggested a rapid turnover in the genes coding for these proteins likely due to a role in prokaryotic innate immunity (Dunin-Horkawicz, Kopec and Lupas, 2014).

WRAP β-propellers belong to the WD40 superfamily, characterised by blades of approximately 40 residues length, carrying a conserved Trp-Asp (WD) sequence motif. This constitutes one of the largest β-propeller superfamilies and corresponds to one of the three major hubs of the β-propeller sequence classification landscape (Chaudhuri, Söding and Lupas, 2008; Kopec and Lupas, 2013; Pereira and Lupas, 2021). The other two are the Asp-Box, characterised by a SxDxGxTW motif (Quistgaard and Thirup, 2009), and the VCBS β-propellers, which contain a conserved cation binding DxDxDG motif and include the β-propellers in α-integrins, various archaeal toxins, and multiple eukaryotic and bacterial lectins (Rigden and Galperin, 2004; Rigden *et al*., 2011; Makarova *et al*., 2019; Pereira and Lupas, 2021). More than 60 unique β-propeller families are classified in ECOD (Cheng *et al*., 2014), most belong to one of these hubs and some form independent clusters that connect to them (Chaudhuri, Söding and Lupas, 2008; Kopec and Lupas, 2013; Pereira and Lupas, 2021). Within these, high sequence and structure symmetry has been also reported for the β-propeller domain in tachylectin-2, a family of VCBS-like eukaryotic and bacterial lectins involved in the innate immune response of arthropods and extensively explored for protein design and the study of β-propeller folding dynamics (Beisel *et al*., 1999; Yadid and Tawfik, 2007, 2011; Vrancken *et al*., 2020; Pereira and Lupas, 2021).

This led us to wonder about the extent of highly repetitive, recently amplified β-propellers in nature. Are WRAP blades especially prone to amplification or is this a more general, inherent property of β-propeller blades? In order to answer this question, we carried out a large-scale search for recently amplified β-propellers belonging to any of the families classified in ECOD.

## Materials and Methods

A summary of the workflow carried out to search, classify, and analyse highly repetitive, putative β-propellers in proteomes across all kingdoms of life is depicted in supplementary figure 1.

### 1. Seed preparation

The searches for highly repetitive β-propellers were carried based on the sequences of individual blades from β-propellers and β-propeller-like folds of known structure. For that, we collected 426 structures of β-propeller-like, β-prisms and β-pinwheels from the ECOD database filtered to 70% sequence identity (ECOD version develop261) (Cheng *et al*., 2014). Their blades were extracted at the structure level by automated detection of internal structure symmetry and structural repeats with CE-Symm 2.1 (Bliven *et al*., 2019). The number of repeat units detected was then compared with the topology annotated in the ECOD database and manually curated.

These β-propeller-like domains were further classified at the sequence level with CLANS (Frickey and Lupas, 2004), which demonstrated that the set collected is overpopulated by the WD40 superfamily. To unskew our dataset and avoid a large number of redundant searches, clusters at a *p*-value of 10^−4^ were automatically detected using the linkage clustering method implemented in CLANS and, for each cluster, a maximum of 3 representatives selected. For that, and for each cluster, their constituent sequences were sorted in descending order of the number of blades and their total coverage of the full domain, and the top ranking 3 β-propellers selected. If a cluster was composed of less than 3 sequences, the full cluster was considered, and if individual sequences did not connect to any other cluster, they were considered a “singleton cluster” and the sequences of their blades saved too. This resulted in 103 clusters, corresponding to an unskewed set of 168 β-propeller-like domains and a total of 1071 blade-like initial sequence seeds.

### 2. Sequence searches

Sequence searches were carried out in parallel for each cluster and individually for each kingdom of life (bacteria, archaea, eukaryotes and virus). By doing so, we avoided that families that were over-represented in a given kingdom (e.g., VCBS sequences in bacteria and WD40 in eukaryotes) took over the PSSMs built and, thus, bias the searches towards that taxonomic group and that family, hindering the detection of under-represented groups. Each search step was composed of three stages: (1) PsiBlast searches for blade-like matches in a sequence database, (2) HMM searches for missed, degenerate blade-like sequences, and (3) selection of continuous, highly repetitive β-propeller-like sequence segments.

In stage (1), the corresponding blade seed sequences of the target cluster were searched with 15 rounds of PsiBlast (Altschul, 1997) against the set of sequences of a given kingdom of life in the NCBI non-redundant sequence database filtered to a maximum sequence identity of 90% (the *nr_bac90, nr_arc90, nr_euk90* or *nr_vir90* databases, respectively) (Zimmermann *et al*., 2018). Only matches at an E-value better than 10^−4^ were considered for PSSM building, and all matches at an E-value better than 1.0 and that covered at least 80% of the query sequence collected as putative blade matches.

At the end of stage (1), the matches collected were used to build an HMM sequence profile, which was used in stage (2) to search for missed blade-like sequences in unusually long matches, in long linkers between two consecutive matches in the same sequence, and N- and C-termini sequence segments. For that, only those matches whose sequence length was at maximum 1 median absolute deviation (MADe) away from the median sequence length in the corresponding set of matches were aligned with MUSCLE (Edgar, 2004) and the resulting alignment trimmed with TrimAl (Capella-Gutierrez, Silla-Martinez and Gabaldon, 2009), removing columns where >70% of the positions were a gap (gap score of 0.30) and sequences that only overlapped with less than 75% of the columns populated by 80% or more of the other sequences. The trimmed alignment was then used to build an HMM profile with HHmake (Söding, 2005), against which all the long matches that were not used to build it were searched. This allowed the detection and further break down of matches that included more than one blade. After that, the set of blades was updated and the HMM rebuilt (following the same procedure) to account for the newly detected matches. Long linkers between two consecutive blade-like matches and with a size compatible with at least one blade were then searched against the updated HMM, the set of blades updated, and the procedure repeated for the N-termini and C-termini of the parental full-length sequence.

In stage (3), the individual blade-like sequence matches detected were grouped into β-propeller-like domains based on the length of the linkers between them. For that, all linkers between two consecutive blade-like matches in the corresponding set of sequences were collected and the median and MADe of their lengths computed. With this, two consecutive blade-like matches were considered to belong to the same β-propeller if the linker between them was less than 3 MADe away from the median linker length. For each putative β-propeller domain with at least 2 consecutive blade matches, their constituent blades were aligned with MUSCLE and the median sequence identity between individual pairs of blades computed. Only those with a median blade sequence identity higher than 60% were considered for further analysis.

### 3. Validation of detected highly repetitive β-propeller-like sequences

In order to validate the boundaries of the blades and make sure that no blade was left behind after stage (3), we further validated our β-propellers with HHrepID (Biegert and Söding, 2008). For that, the full-length sequences encompassing the selected β-propellers were analysed, searching for significant repeat units at a *p*-value of 10^−1^. For a given target full-length sequence, if the repeats detected with HHrepID covered more of it than the putative blades already annotated in the previous steps, this region was further compared with the previous annotations.

As the host full-length protein sequence may contain other repeat patterns in addition to those in β-propeller regions, and HHrepID is able to differentiate between different types of repeat units, only those repeat types in regions matching already annotated blades were considered. However, even if a given repeat type can be assigned to correspond to a β-propeller region, we cannot assume that the repeat unit corresponds to a blade; it can also correspond to a larger unit or even to a full-length β-propeller depending on the evolutionary history of the target protein. Thus, and to detect such cases, the number of blades within a given repeat unit was always predicted based on the median length of the blades of the corresponding annotated β-propeller. If a given repeat unit could accommodate at least 2 blades, it was further analysed with HHrepID iteratively until no more repeats with a blade-like size were detected. On the other hand, if a repeat unit could accommodate one blade only, it was considered a putative blade.

The intervals of putative blade-like repeat units and sequence-based blade like matches were compared and merged to maximize the length of the detected β-propeller. If a given repeat type annotated with HHrepID encompassed more than one of the β-propellers annotated before, these were merged into one β-propeller, whose boundaries were defined to maximise the coverage of the full-length protein. On the other hand, if only one β-propeller would fit into the repeated region, the interval with the largest number of repeats/blades was considered the β-propeller. This procedure allowed for the detection of further degenerate blades and the merging of seemingly short β-propeller-like segments, into longer putative β-propeller domains. These β-propellers were again analysed for the median sequence identity of their individual blades, resulting in 10,480 putative highly symmetric β-propeller domain sequences with a median blade sequence identity higher than 60%, from a total of 9,919 protein sequences.

### 4. Sequence annotation

The putative highly-repetitive β-propeller domains identified were assigned an ECOD family with HHsearch (Söding, 2005). For that, each β-propeller sequence was searched against a database of HMM profiles built for the ECOD database filtered to 70% maximum sequence identity (the HHpred ECOD70 database as of November 2021, (Zimmermann *et al*., 2018)) and assigned the best match at a probability better than 50%.

To analyse the domain environments of each β-propeller, i.e., annotate the domain composition of the proteins bearing the highly repetitive β-propellers to classify them into “globally” or “locally” repetitive, full-length sequences were annotated similarly (using the HHpred database as of May 2020), but iteratively, so that sequence regions not yet mapped to a domain in the previous iteration were searched individually again. A maximum of five iterations were carried out and only the best matches outside already annotated β-propeller regions at a probability better than 50%, larger than 40 residues and that covered at least 30% of the target profile considered.

### 5. Three-dimensional structure prediction and model quality estimation

Three-dimensional structural models of selected repetitive β-propeller examples were produced with AlphaFold v2.1.2 monomer (Jumper *et al*., 2021). Modelling was carried out with af2@scicore v2.2.0 (https://git.scicore.unibas.ch/schwede/af2-at-scicore) using default settings. For each example, five models were generated. Model quality was estimated based on the predicted LDDT scores reported by AlphaFold but also independently, with QMEANDisCo, through the SWISS-MODEL “Structure Assessment” tool (Waterhouse *et al*., 2018; Studer *et al*., 2020).

### 6. Identification of non-coding β-propeller fragments

Given that most of the highly repetitive β-propeller regions were found at protein extremities (N- or C-terminal regions), we analysed the immediately adjacent non-coding genomic regions that flank their corresponding genes for nucleotide segments with β-propeller-encoding potential. For that, the genome assembly corresponding to each protein was mapped using Biopython.Entrez (Cock *et al*., 2009) by first querying for each unique EntrezID the corresponding NCBI Identical Protein Group (IPG), collecting the accession codes of all genome assemblies matched and then filtering out repeated assemblies (i.e., GCA assemblies for which there is an identical, GCF, entry on RefSeq).

For all protein coding regions mapped, the nucleotide sequence of the immediate 5’ and 3’ non-coding intergenic regions (i.e., the genomic regions between the current open reading frame (ORF) and the two closest, flanking ORFs excluding pseudogenes, in the same direction) were collected and translated in all three frames possible in the same direction of the target ORF. Stops resulting from stop codons were replaced by X, emulating an unknown amino acid. These protein-like sequences were then iteratively annotated against the ECOD70 database using the same approach described above; for this case, a maximum of five iterations were carried out and only the best matches to β-propeller-like folds at a probability better than 1%, larger than 20 residues and with a maximum 15% content of stop codons were considered.

## Results

We started with the blades of 168 non-redundant β-propellers and β-propeller-like domains of known structure, and used them as seeds for deep sequence similarity searches in the non-redundant sequence database at NCBI (*nr_bac, nr_arc, nr_euk* and *nr_vir;* Fig. S1) (Zimmermann *et al*., 2018). This initial set comprised not only the β-propeller domains classified in ECOD but also β-pinwheels and type II β-prisms, which are repetitive, symmetric folds based on the repetition of a 4-stranded blade-like unit distantly related to β-propeller blades (Kopec and Lupas, 2013). β-pinwheels and type II β-prisms are represented by the DNA-binding domains (Corbett, Shultzaberger and Berger, 2004) of bacterial type IIA topoisomerases and *Galanthus nivalis* agglutinin-related lectins (GNA-related lectins) (Hester and Wright, 1996; Kaus *et al*., 2019), respectively.

In our searches, we specifically probed for those sequences with at least two consecutive non-overlapping matches to the seed blade and a median internal blade-to-blade sequence identity higher than 60%. We selected this threshold as, apart from WRAP, no natural β-propeller was known with such a high internal sequence symmetry (Pereira and Lupas, 2021). Our approach resulted in 10,474 putative β-propeller domains across 9,919 host protein sequences from all kingdoms of life.

### 1. β-propeller families with highly repetitive β-propellers

HHsearches against the ECOD database allowed us to map each of these domains to 32 out of 65 unique ECOD β-propeller families (Fig. 1a). While a 50% probability threshold was used for annotation, most matches have a probability better than 90% (Fig. 1b). Most of these correspond to those of β-propeller domains usually involved in macromolecular interactions, but mappings to at least eight commonly associated with a catalytic activity were also found. Still, only 17 sequences did not find significant matches to any family (UNKNOWN label), while 3 found their best match in a non-propeller fold (NO_PROPELLER label); these 3 cases were excluded from further analysis.

**Figure 1.**
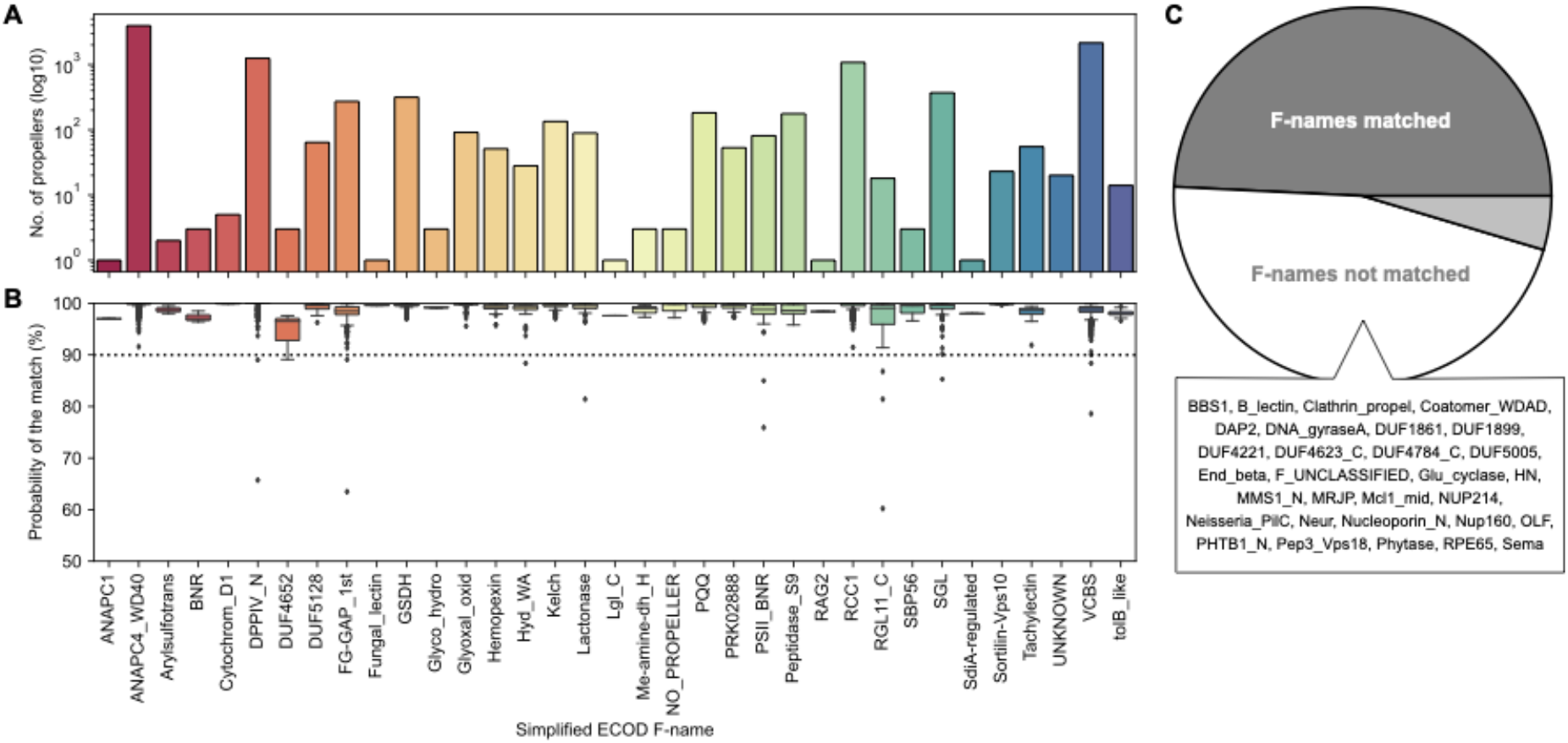
The ECOD families matched. (A) Frequency of the different ECOD F-names (in their simplified form) matched. (B) Boxplots of the HHsearch probability of the match, per family. (C) Proportion of ECOD F-names mapped, with a list of those without any match to β-propeller-like sequences in our dataset. The UNKNOWN family corresponds to the sequences without significant matches to any of the 65 ECOD F-names.

The sequences in our dataset span a wide range of median sequence identities of their blades, reaching up to full identity in half the lineages of the dataset (Fig. S2a).

Half of the sequences collected are of the WD40 superfamily, to which WRAP belongs. The WRAP domain in Npun_R6612 of *Nostoc punctiforme* PCC 73102 was the β-propeller with the highest internal sequence symmetry we had found up to that point and contains no synonymous substitutions at the DNA level (Fig. 2), pointing to a very recent amplification event (Dunin-Horkawicz, Kopec and Lupas, 2014). Keeping in mind our hypothesis that new β-propellers continuously arise through cycles of amplification and differentiation, we attempted to track the process of differentiation in this very recently arisen β-propeller through the emergence of synonymous mutations in close homologs. Indeed, these homologs show a proportion of synonymous mutations, along with a higher number of non-synonymous ones, but to our surprise appeared to have resulted from independent amplification events, as judged by the presence of positions at which all the blades of one β-propeller contained a different nucleotide than all the others (Figs. 2 and S3). Thus, even at pairwise sequence identities above 80%, ungapped, these β-propellers are only homologous at the level of the blade that had been amplified, but analogous in the fully amplified form.

**Figure 2.**
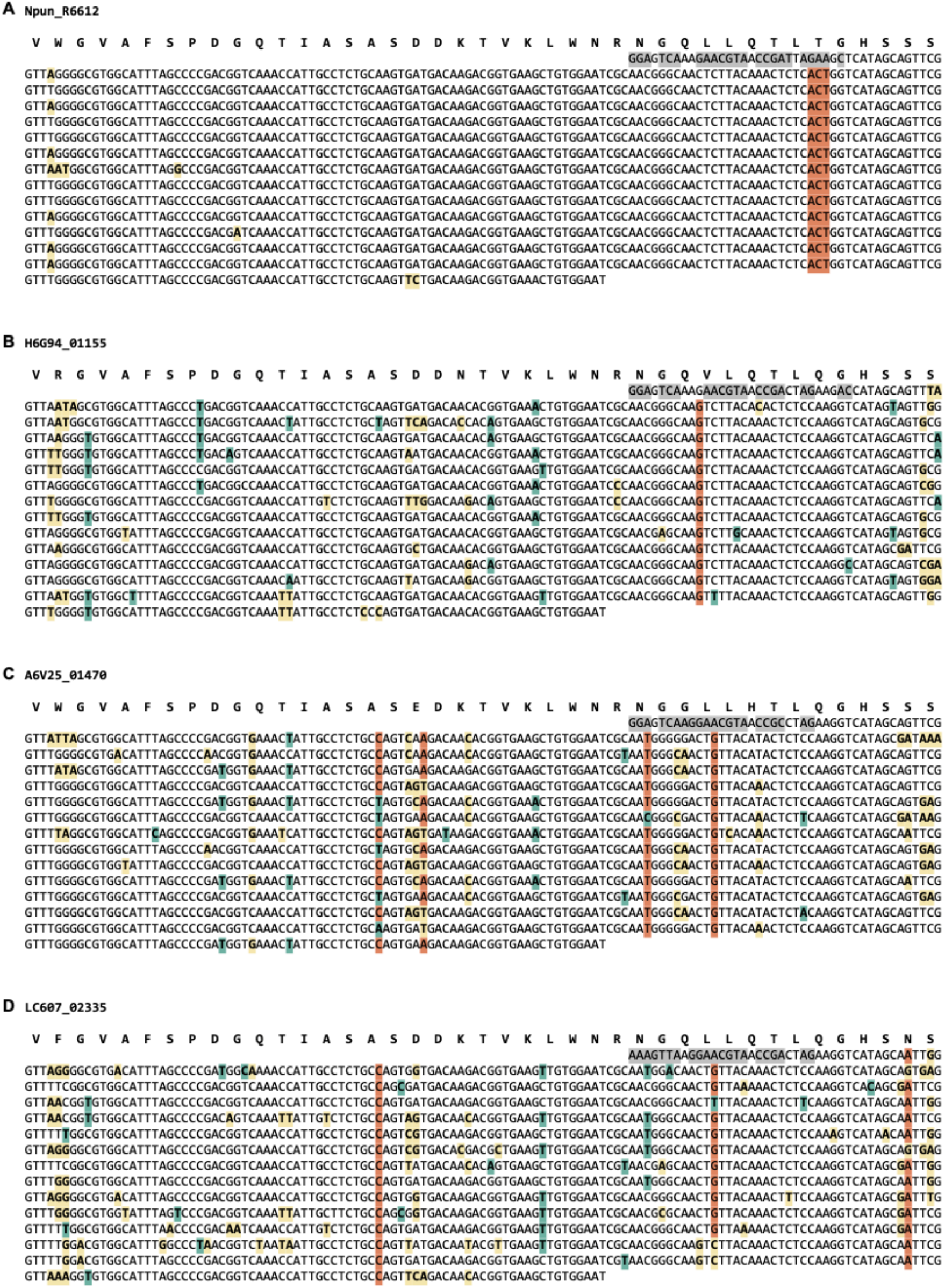
Synonymous and non-synonymous mutations in the nucleotide sequence of four closely related WRAP domains. Synonymous mutations relative to the majority rule consensus of the respective β-propeller are highlighted in teal, and non-synonymous ones in yellow. Positions where a nucleotide is present in at least two thirds of the blades of one β-propeller, but different from the equivalent nucleotides in at least two of the other β-propellers are highlighted in orange. The nucleotide sequence encoding the first β-strand of the structure is different from that of the equivalent strand in the amplified region and highly conserved between homologs, and is highlighted in grey. (a) Npun_R6612 of *Nostoc punctiforme* PCC 73102 (ACC84870.1), (b) H6G94_01155 of *Nostoc punctiforme* FACHB-252 (MBD2609895.1), (c) A6V25_01470 of *Nostoc sp*. ATCC 53789 (RCJ36011.1), and (c) LC607_02335 of *Nostoc sp*. CHAB 5824 (MCC5641816.1). These proteins have at least 80% pairwise sequence identity (Fig. S3).

The VCBS superfamily is the second most sampled, encompassing about 20% of the β-propeller-like sequences in the dataset. We previously demonstrated that VCBS β-propellers form a large and diverse hub in the β-propeller classification landscape that is tightly connected to the FG-GAP/α-integrin, tachylectin-2 and Family 11 rhamnogalacturonan lyase families and is bridged to the WD40 supercluster by the PQQ family (Pereira and Lupas, 2021). In line with this, we also found highly repetitive β-propeller-like sequences assigned to the FG-GAP/α-integrin (2%), tachylectin-2 (0.7%), Family 11 rhamnogalacturonan lyase (RGL11, 1%) and PQQ (1.7%) families. While VCBS-like β-propellers are typically associated with the recognition of different carbohydrates (but also protein-protein interactions) (Pereira and Lupas, 2021), RGL11 β-propellers are catalytic and involved in the cleavage of the rhamnogalacturonan type-I region of plant cell wall pectin (Ochiai *et al*., 2007, 2009), and PQQ β-propellers in protein binding in various biological processes, including outer membrane protein assembly, alcohol metabolism and unfolded protein response (Ghosh *et al*., 1995; Kim and Paetzel, 2011; Kopec and Lupas, 2013).

The third most populated group, making up 10% of the set, is RCC1/BLIPII. Well-known members of this group are the β-propeller domains in the eukaryotic Regulator of Chromosome Condensation 1 (RCC1) protein, a nuclear interactor involved in the regulation of spindle formation and nuclear assembly during mitosis (Hadjebi *et al*., 2008; Stevens and Paoli, 2008), and the bacterial β-lactamase inhibitor protein II (BLIP-II), a secreted binder produced by soil bacteria as a potent β-lactamase inhibitor (Park and Lee, 1998; Brown *et al*., 2013). In our dataset, members of this group are mostly of bacterial origin, but a few eukaryotic and viral members were also collected (Fig. 3b).

**Figure 3.**
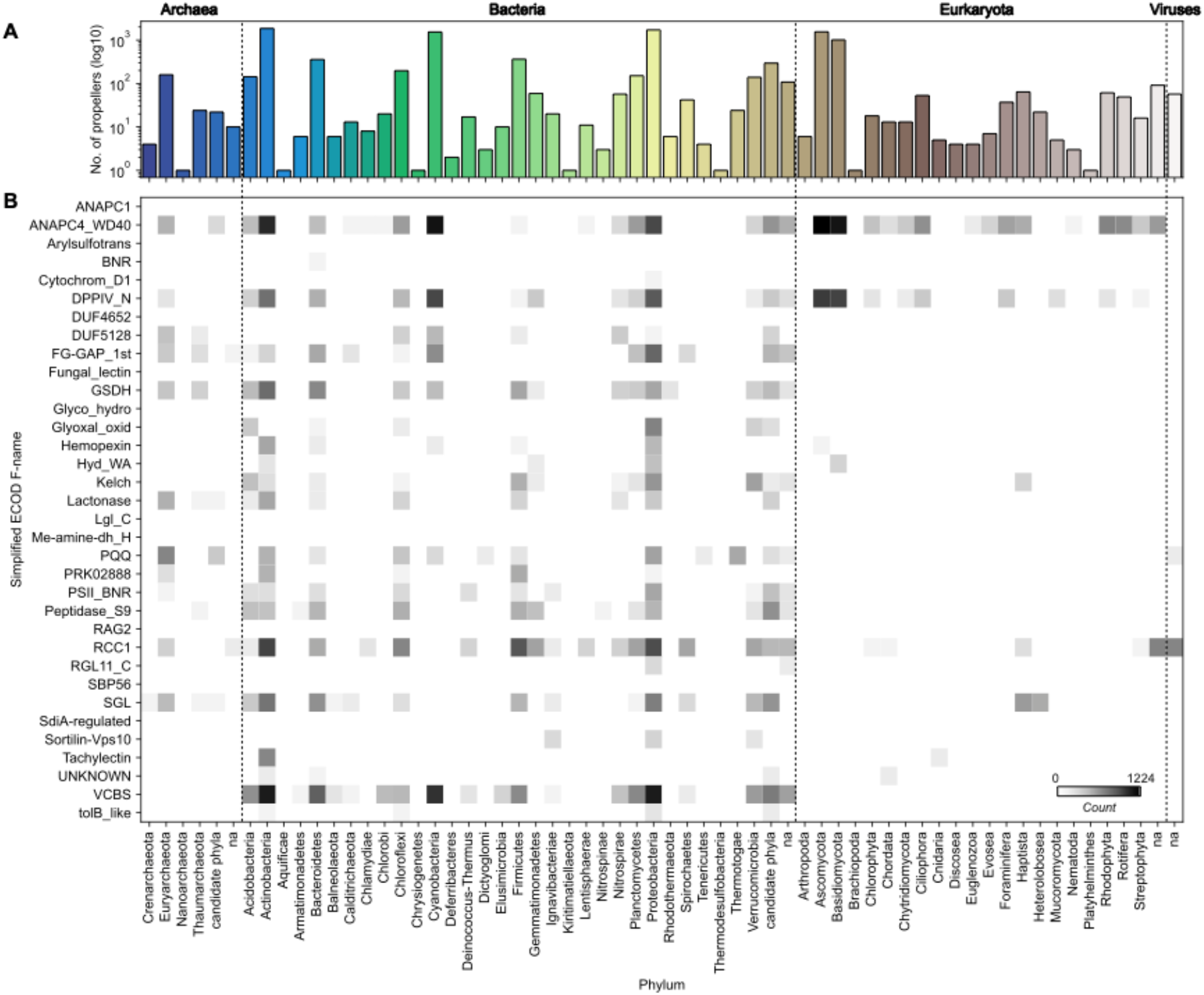
Taxonomic distribution of highly repetitive β-propeller sequences. (A) The frequency, in logarithmic scale, of highly repetitive β-propeller sequences from different phyla in the collected dataset. (B) Heatmap of the frequency of highly repetitive β-propeller sequences of different ECOD families in different phyla. Colors are in logarithmic scale.

The fourth largest set of sequences maps to the Asp-Box superfamily (e.g., Sortillin-, PSII- and SGL-like β-propellers), making up to 5% of the dataset. The remaining sequences belong to, among others, (1) the DPPIV and Peptidase S9 families (Pfam: PF00930 and PF02897), found in multiple bacterial and eukaryotic serine, prolyl oligopeptidases where the β-propeller domain controls the access to the active site (Fülöp, Böcskei and Polgár, 1998; Polgár, 2002; Hiramatsu *et al*., 2003), (2) the Hemopexin family (Pfam: PF00045), which encompasses the domains in various exported eukaryotic proteins involved in haem transport and protein-protein interactions (Das *et al*., 2003), (3) the Glucose/Sorbosone dehydrogenase (GSDH) family (Pfam: PF07995), which are bacterial pyrroloquinoline quinone (PQQ)-dependent enzymes (Oubrie *et al*., 1999), (4) two distinct Lactonase-like families (Pfam: PF10282), that bind the co-factors of various enzymes involved in the breakage of lactone rings (Amara *et al*., 2011) and include the protein-binding domain of the archaeal Lp49 surface antigen (Giuseppe *et al*., 2008), and (5) the Kelch superfamily (Pfam: PF01344), whose members are functionally diverse and may be involved in protein binding (Gupta and Beggs, 2014).

On the other hand, and although they were included in the initial set of seed sequences, no highly repetitive members were found for families with a β-prism or a β-pinwheel fold, nor for members of the Ph1500 family, which form, by oligomerization, the largest, 12-bladed β-propellers known to date (Varnay, 2009).

### 2. Taxonomic distribution of proteins with highly repetitive β-propellers

Within the β-propeller-like sequences in our dataset, we found representatives from all kingdoms of life, including viruses (Fig. 3). While examples for most phyla could be found, the most prevalent are Actinobacteria, Cyanobacteria, Proteobacteria, Ascomycota and Basidiomycota, followed by Bacteroidetes, Chloroflexi, Firmicutes and Euryarchaeota (Fig. 3a). Indeed, ∼70% of the β-propellers collected (and the same proportion of full-length host protein sequences) belong to bacterial species, and it is in bacteria that the largest variety is found, followed by archaea, eukaryotes, and viruses (Fig. 3b). Sequences from Ascomycota and Basidiomycota, two phyla of fungi, make ∼25% of the dataset and fully 84% of the eukaryotic sequences.

When looking at the individual taxonomic distribution of specific families, there are families that are widespread while others have highly repetitive members in just a few taxonomic groups (Fig. 3b). For example, among highly repetitive β-propellers, WD40 members are widespread, but especially prevalent in fungi, Actinobacteria, Cyanobacteria and Proteobacteria, VCBS members are only found in bacteria, FG-GAP/α-integrin members are found in both archaea and bacteria, and tachylectin-like members are unique to Actinobacteria and, surprisingly, Cnidaria.

### 3. The diverse blade numbers of highly repetitive β-propellers

Sequences in our dataset may contain as few as 2 or as many as 47 highly similar consecutive blades (Fig. 4). Their number depends primarily on the β-propeller family (Figs. S2b and S4). For example, highly repetitive WD40-like β-propellers have a median number of 7 blades, while VCBS-like have 6 and DPPIV-like 13. Looking at the distribution of blade numbers in individual lineages, we observe that there is a continuum of blade numbers, which are however biased towards the median number in their family. For example, for β-propellers whose best match in ECOD is a WD40 β-propeller, the histogram of the number of consecutive highly identical blades has peaks at 7 and 14, with a continuum of less frequent intermediate blade numbers.

**Figure 4.**
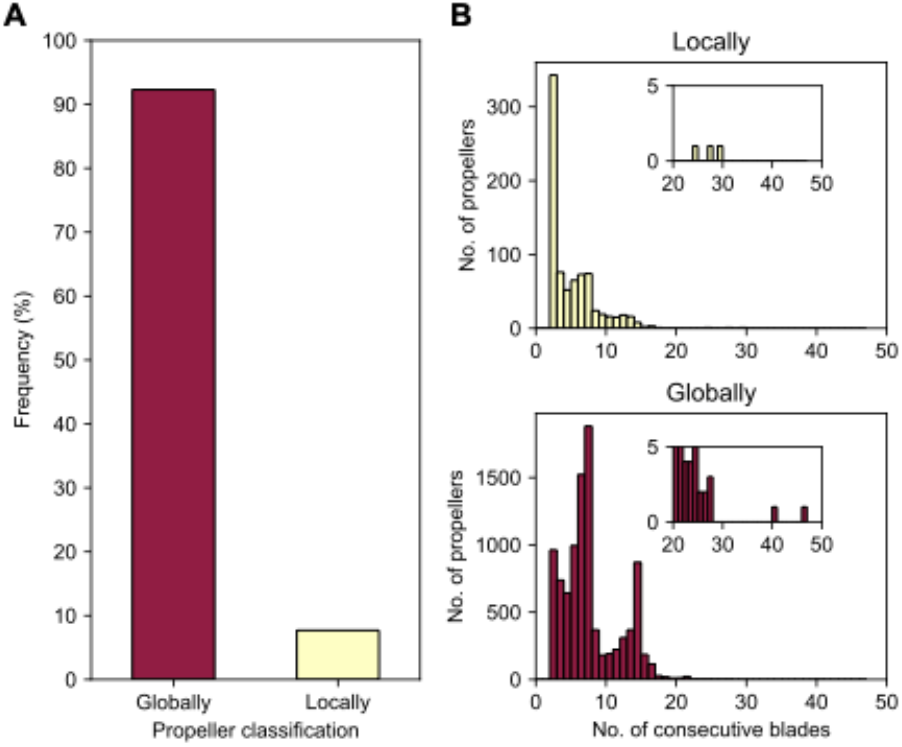
Globally versus locally highly repetitive β-propellers. (a) The proportion of β-propellers in the dataset categorised into “globally” or “locally” based on the absence or presence of non-repetitive β-propeller matches in their immediate domain environment. (b) Histograms for the absolute number of consecutive highly identical blades, separated by highly repetitive β-propeller type.

However, HHsearches with the full-length sequences of all 9,919 proteins containing highly repetitive β-propellers indicated that in 8% of the cases these regions have adjacent, non-highly repetitive β-propeller matches, suggesting they correspond to parts of larger, degenerate β-propellers in which only some consecutive blades are highly similar. Such cases we denote as “locally repetitive β-propellers”. The remaining 92% correspond to continuous regions of consecutively amplified, highly similar blades. These correspond to “globally repetitive β-propellers” (Fig. 4a). There are no family or taxonomic differences between globally and locally repetitive β-propellers, yet their preferences for the number of highly similar blades differ (Fig. 4b).

#### 3.1. The blade numbers of locally repetitive β-propellers

The most frequent number of highly similar blades in locally repetitive β-propellers is 2, but, in some instances, it may be as high as 29 (Fig. 4). A representative example for a locally repetitive β-propeller is the β-propeller domain of the inactive dipeptidyl peptidase 10 (DPPY) of a wild silk moth (*Bombyx mandarina*) (XP_028029463.1, Fig. 5). This 929-residue membrane protein contains 7 divergent β-propeller blades of the DPPIV family with a median sequence identity of 14%, followed by 2 blades with a pairwise sequence identity of 100% (Fig. 5a) that resulted from the amplification of a single exon. While DPPY β-propellers adopt an 8-bladed fold (Bezerra *et al*., 2015), we obtained two possible models for the 9 blades of the *Bombyx* protein (Figs. 5c,d). While an early version of ColabFold (Mirdita, Ovchinnikov and Steinegger, 2021) produced a 9-bladed β-propeller (Figs. 5c), recent versions of ColabFold and AlphaFold (Jumper *et al*., 2021) (as of February 2022) always produced 8-bladed β-propellers, where the first identical blade is part of the domain and the second identical blade is modelled as a single, unpacked blade (Figs. 5d).

**Figure 5.**
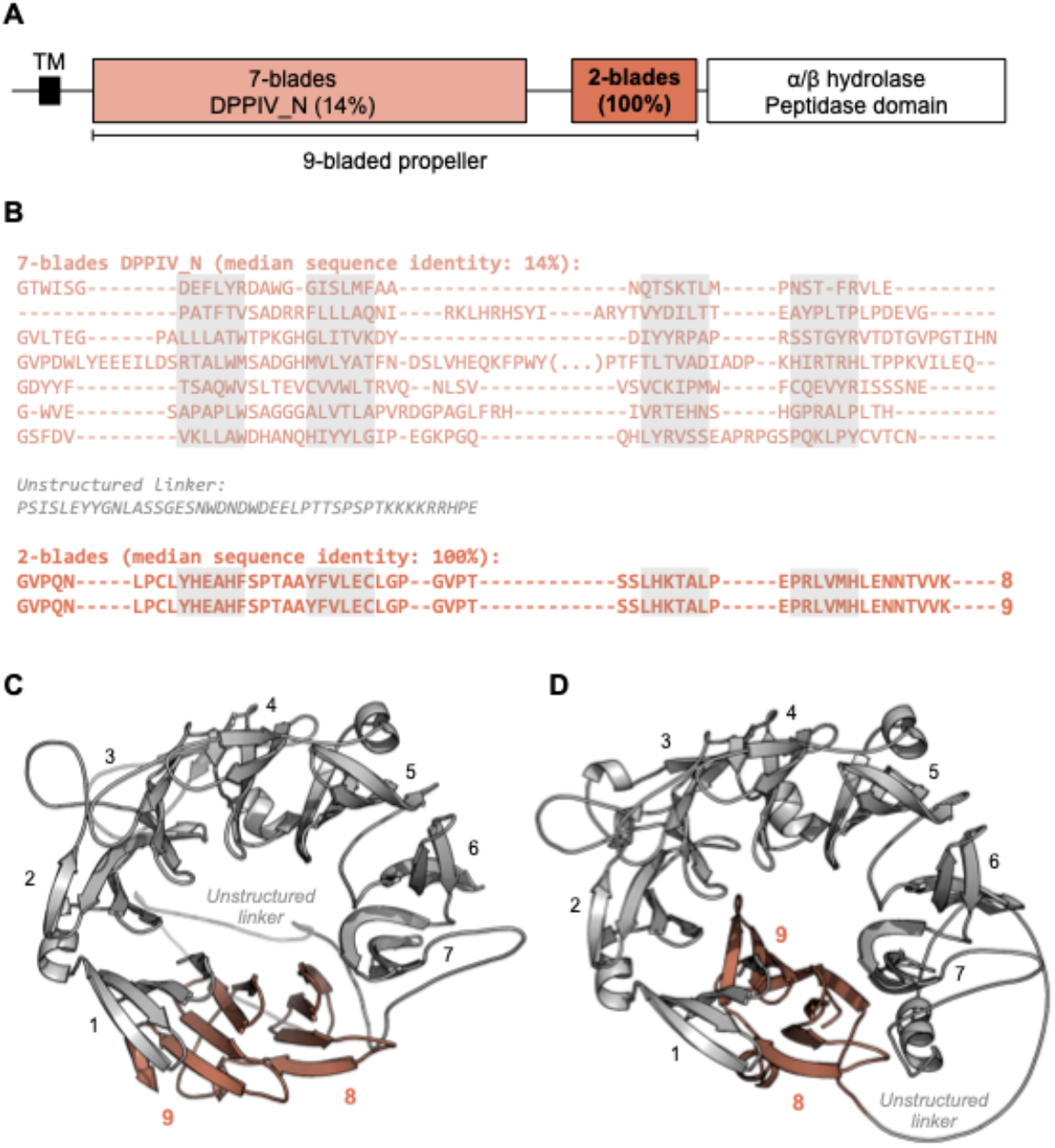
The locally highly repetitive β-propeller of *Bombyx mandarina* inactive dipeptidyl peptidase 10 (DPPY) (XP_028029463.1). (A) Predicted domain composition of the full-length DPPY sequence. (B) Annotated β-propeller sequence, highlighting the two highly identical, consecutive blades. β-propeller blade sequences were aligned with Promals3D (Pei and Grishin, 2014) based on their predicted structure. The individual strands in each β-propeller blade are highlighted in grey. (C) Three-dimensional model of the β-propeller region, predicted with ColabFold MMseqs protocol as of August 2021 (Mirdita, Ovchinnikov and Steinegger, 2021) (QMEANDisCo Global (Studer *et al*., 2020) score of 0.59±0.05, average pLDDT (Jumper *et al*., 2021) of 82.9). (D) Three-dimensional model of the β-propeller region, predicted with AlphaFold v2.1.2 (Jumper *et al*., 2021) (QMEANDisCo Global (Studer *et al*., 2020) score of 0.62±0.05, average pLDDT (Jumper *et al*., 2021) of 85.3).

#### 3.2. The blade numbers of globally repetitive β-propellers

In contrast to the locally repetitive β-propellers, globally repetitive ones strongly prefer larger numbers of blades (Figs. 4b-d and S4). In fact, they often have a greater number of blades than the median number for their lineage. For example, the β-propeller region in the WD40 repeat-like protein KAE9403162.1 from *Gymnopus androsaceus* JB14 corresponds to 15 WD40-like consecutive blades with a median sequence identity of 83% (Fig. 6a). AlphaFold predicts, at high confidence (average pLDDT of 89%), that this region folds into two individual β-propeller domains, one with 7 blades and the other with 8 (Fig. 6b).

**Figure 6.**
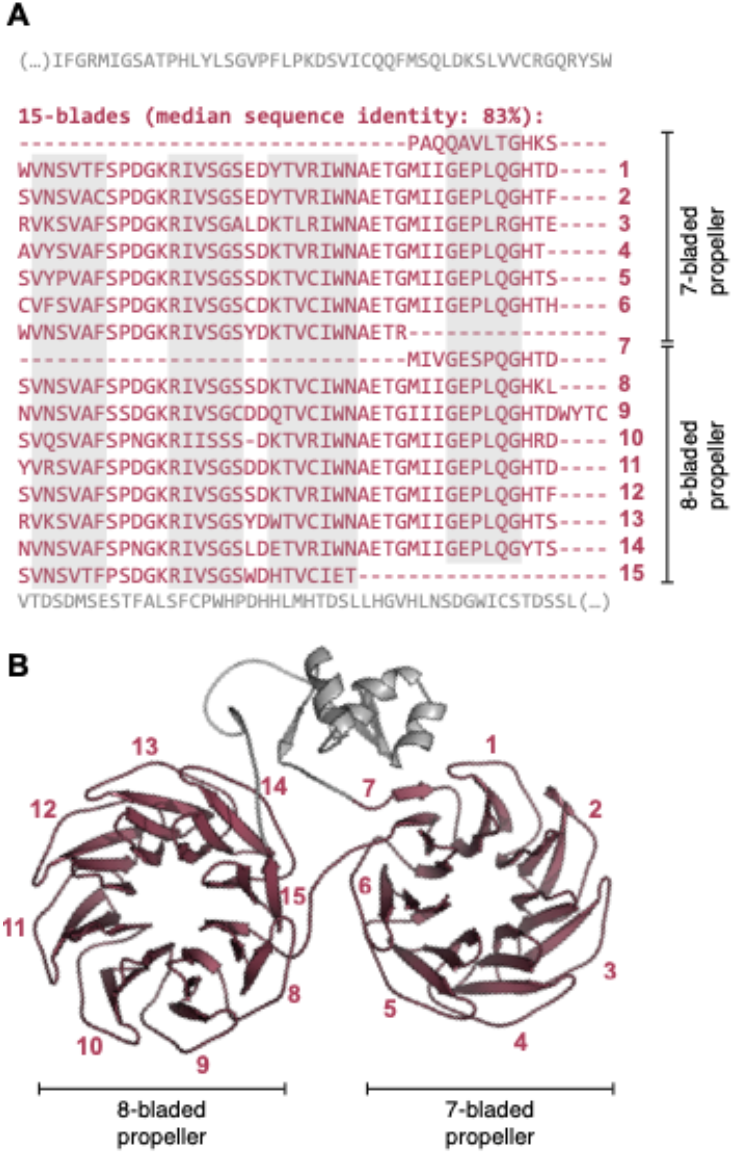
The globally highly repetitive β-propeller of WD40 repeat-like protein KAE9403162.1 from *Gymnopus androsaceus* JB14. (A) Annotated β-propeller sequence, highlighting the 15 nearly identical, consecutive blades and the flanking non-propeller sequences. The individual strands in each β-propeller blade are highlighted in grey. (B) Three-dimensional model of the β-propeller and immediate flanking regions predicted with AlphaFold v2.1.2 (Jumper *et al*., 2021) (QMEANDisCo Global (Studer *et al*., 2020) score of 0.65±0.05, average pLDDT (Jumper *et al*., 2021) of 88.9). Modelling was carried out with the full-length protein sequence but only the region whose sequence is shown in (A) is depicted.

However, modelling globally repetitive β-propellers is not always so straightforward. For example, the highest number of blades in our set is 47, found in a hypothetical NACHT ATPase from *Penicillium camemberti* (CRL31138.1) (Fig. S5a). The 47 blades are compatible with multiple combinations of 6 to 9-bladed β-propellers, but no high-quality three-dimensional model could be obtained through AlphaFold (Fig. S5b). In all 5 models generated, β-propeller silhouettes can be seen but they all collapse into each other. A reason may lie in their extreme level of sequence symmetry: while the median blade identity is 81%, consecutive fragments of 2 blades have a median sequence identity of 100% at both the protein and the nucleic acid levels. The unit of repetition is thus 2-bladed and the sequence is extremely symmetric (Fig. S5c), representing the only case of amplification from a 2-bladed fragment we have observed so far. This makes it difficult to generate meaningful multiple sequence alignments for protein structure prediction based on co-evolutionary potentials and impossible, so far, to predict how many β-propellers it may be able to fold into. Its complete internal sequence identity, even at the nucleotide level, also makes it difficult to judge whether there are indeed 47 blades, given the problems of assembling different fragments with identical overlaps. Even though not as pronounced, this may be a recurrent problem in our highly repetitive β-propellers.

We also found globally repetitive β-propellers with fewer blades than the median number for their lineage and interpret these as fragments of full domains (Fig. S4). One example is provided by the 2 blades found in the hypothetical protein DL7770_005219 from *Menosporascus sp*. CRB-9-2 (Fig. 7). This protein carries two WD40-like tandem blades with a sequence identity of 92%, which are flanked by an N-terminal α-helical region and a C-terminal region rich in β-strands. We obtained different models for this part of the protein (Figs. 7b-d) and, in all cases, the two highly similar blades are predicted as expected, but in each model the C-terminal region appears to provide additional blade-like supersecondary structures. Indeed, HHsearches with just the C-terminal region showed significant matches to segments of 3 consecutive WD40-like blades. This suggests that this region may correspond to a disrupted β-propeller that is quickly diverging, similarly to those described in a previous work for WRAP homologs (Dunin-Horkawicz, Kopec and Lupas, 2014).

**Figure 7.**
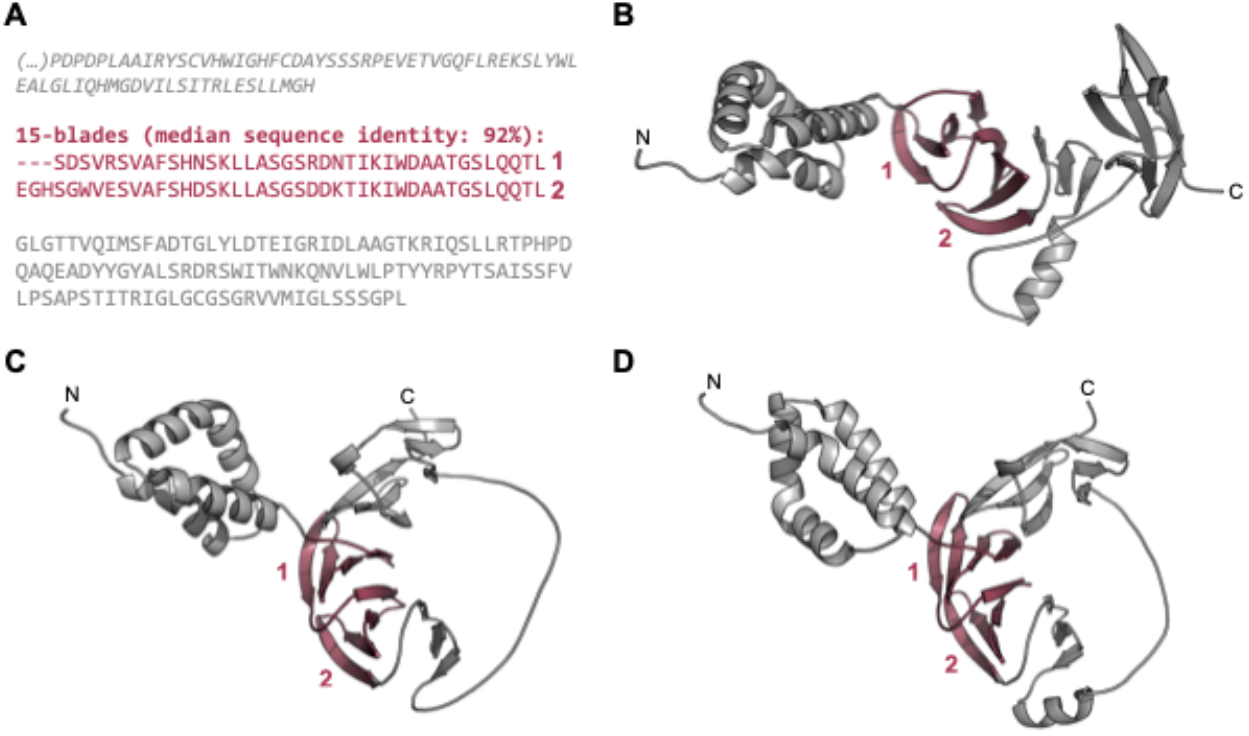
The globally highly repetitive β-propeller of *Menosporascus sp*. CRB-9-2 hypothetical protein DL7770_005219 (RYP84315.1). (A) Annotated β-propeller sequence, highlighting the two highly identical, consecutive blades and the flanking non-propeller sequences. The individual strands in each β-propeller blade are highlighted in grey. (B-D) Three-dimensional models of the β-propeller and immediate flanking regions predicted with AlphaFold v2.1.2 (Jumper *et al*., 2021). Modelling was carried out with the full-length protein sequence but only the region whose sequence is shown in (A) are depicted. (B) The top ranking model (QMEANDisCo Global (Studer *et al*., 2020) score of 0.57±0.05, average pLDDT (Jumper *et al*., 2021) of 80.1), (C) the second best model (QMEANDisCo Global score of 0.58±0.05, average pLDDT of 75.9), and (D) the third best model (QMEANDisCo Global score of 0.58±0.05, average pLDDT of 71.0).

### 4. The evidence for disrupted β-propellers

While it is possible that globally repetitive β-propeller fragments may occur within a non-propeller context, most of the sequences we found are located less than 50 residues away from one of the termini of their full-length protein, more often the C- than the N-terminus (Fig. 8). Given earlier evidence of highly repetitive WRAP-like β-propellers disrupted by either frameshifts or in-frame stop codons, and the overall continuous distribution of blade numbers in our dataset, we therefore searched for putative β-propeller blades encoded in the 5’ or 3’ regions as evidence for disrupted genes.

**Figure 8.**
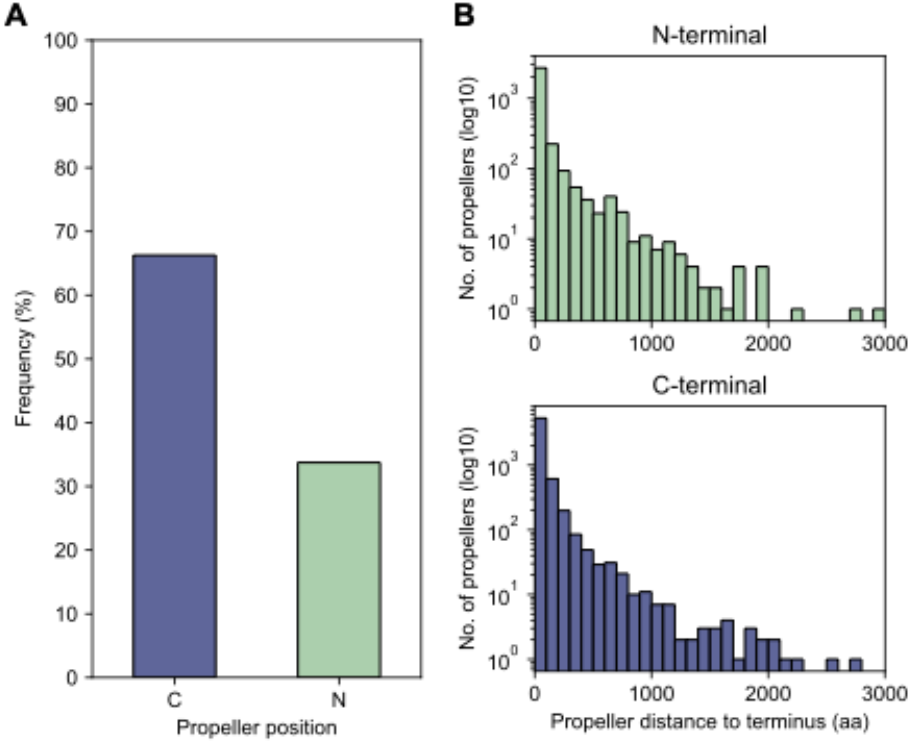
The proximity of globally highly repetitive β-propellers to the termini. (a) The proportion of β-propellers in the dataset localised close to one of the termini of the corresponding host protein. For each globally repetitive β-propeller in the dataset, the distance to both the N- and C-termini was computed and the closest chosen as the closest to the β-propeller position. (b) Histogram of the number of amino acids separating the β-propeller from its closest terminus.

We could map 7329 (76%) globally repetitive β-propellers to 3,175 unique genome assemblies and found putative β-propeller fragments in the neighbourhood of 21% of these (corresponding to 1,553 globally repetitive β-propellers across 584 assemblies). While 83% of the fragments are of low confidence (Fig. 9a), as the probability of their HHsearch match is lower than 50%, the probability distribution is bimodal and 13% (corresponding to 428 fragments close to 320 globally repetitive β-propellers) have a HHsearch probability above 70%. High confidence fragments tend to be closer to the gene containing the globally repetitive β-propellers, which per se are closer to the terminus (Figs. 9b,c). There is no clear family or taxonomic preference for such putatively disrupted β-propellers (Fig. S6).

**Figure 9.**
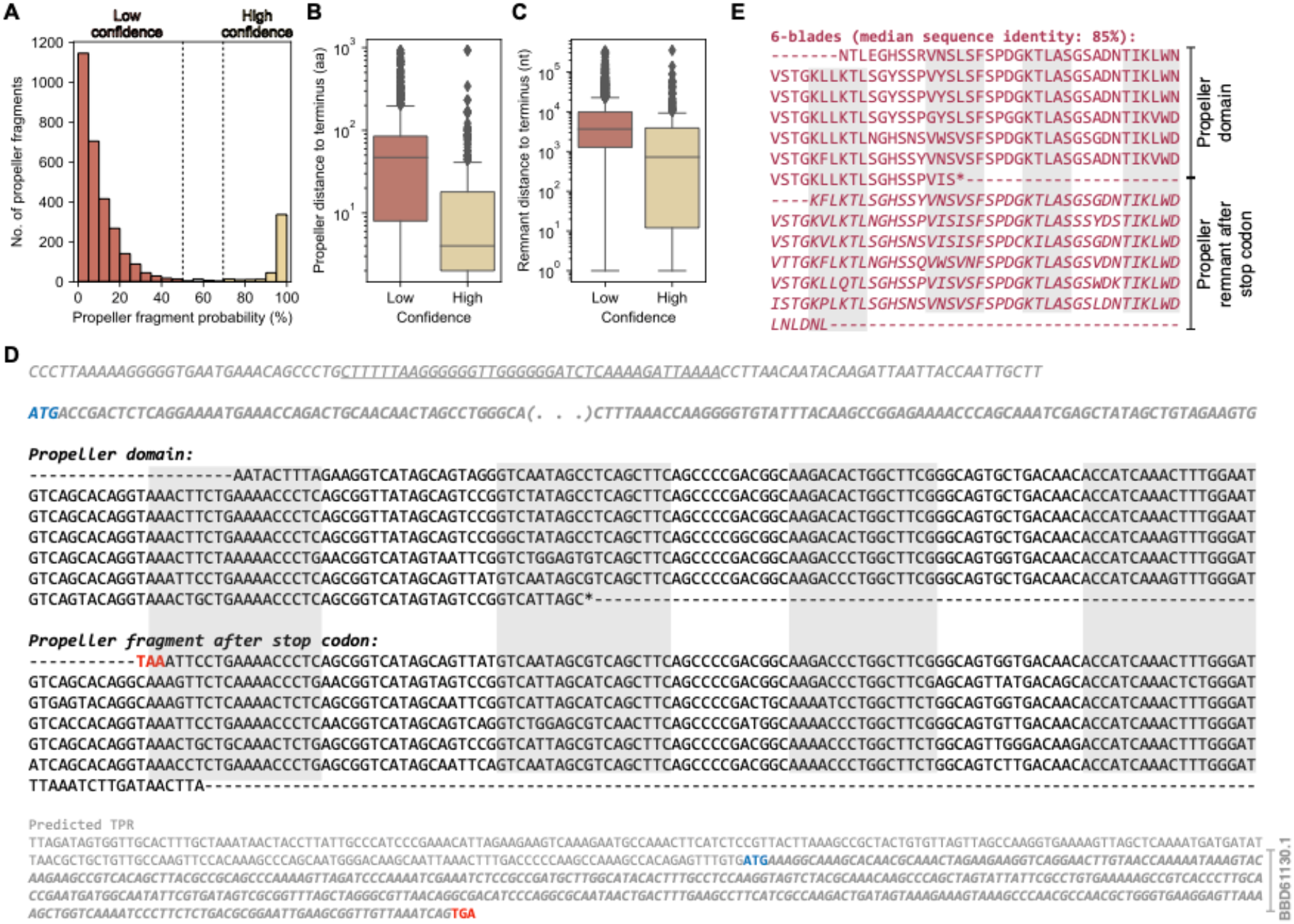
The β-propeller fragments found in the genomic neighbourhood of globally highly repetitive β-propellers. (a) Histogram of the HHsearch probability of the identified putative fragments. Boxplots for the distance of the (b) highly repetitive β-propeller (in amino acids) and (c) corresponding putative β-propeller fragment (in nucleotides) to the closest host protein/gene terminus, separated by confidence level. (d) Nucleotide sequence and genomic context of the exemplary globally repetitive β-propeller from *Nostoc sp. HK-1* hypothetical WD40 repeat-containing protein BBD61132.1. The starting codon is highlighted in blue and in frame, disruptive, stop codons in red. The nucleotide sequence coding for the N-terminal region of the protein is shortened and the regions coding for the strands making up the different blades are highlighted in grey. (e) Amino acid sequence of the 6 highly similar blades annotated at the C-terminus of BBD61132.1 plus the additional 6 found beyond the stop codon (and that make the β-propeller fragment).

One high-confidence example is shown in figure 9d. It is exactly downstream from the open reading frame (ORF) for *Nostoc sp. HK-1* hypothetical WD40 repeat-containing protein BBD61132.1, which contains 6 WD40-like blades with a median sequence identity of 85% (Fig. 9e). Aligning the fragment nucleotide sequence with that coding the β-propeller in BBD61132.1 revealed that it is the result of a deletion that placed a premature stop codon in frame. Right after this stop codon there is an out-of-frame ORF with six other blades (the fragment), which are not only highly similar within themselves, but also to those in BBD61132.1, and a C-terminal TPR repeat that is partially assigned to ORF BBD61130.1. It is thus likely that BBD61132.1 was a longer protein that contained a C-terminal TPR repeat and an at least 12-bladed globally repetitive β-propeller that was disrupted.

Another example is shown in figure S7. In this case, a hypothetical protein, DMF28_08825 (PYL67592.1) from an unknown Verrucomicrobia bacterium, contains a globally repetitive β-propeller with 4 Kelch-like blades of median sequence identity of 76%, followed by a segment without similarity to any known protein (Fig. S7a). The analysis of the nucleotide sequence shows that there was most likely a large deletion in the fifth blade, which moved the gene out of register and caused a sixth blade to be encoded in another frame, hence its amino acid sequence not matching anything in the protein database. Following an in-frame stop codon, there is a further seventh blade and the part required to complete the first blade (Figs. S7b,c). We conclude that this protein used to have a 7-bladed Kelch-like β-propeller, which is now disrupted by a frameshift and a stop codon.

Our analysis provides a lower bound for disrupted β-propellers, as we note that we did not identify additional blade sequences in the vicinity of every β-propeller we think is disrupted, such as in the case as the highly repetitive β-propeller in the hypothetical protein DL7770_005219 described above.

## Discussion

The range of internal sequence symmetries in β-propellers has led us to propose that their different lineages arose by independent cycles of amplification and differentiation (Chaudhuri, Söding and Lupas, 2008). Here we analysed whether these cycles are still ongoing within individual lineages by tracking the occurrence of highly symmetrical β-propellers. We find that β-propellers with a median internal sequence identity of more than 60% are widespread across all major β-propeller lineages, including the three superclusters formed by WD40, VCBS and Asp-Box proteins, respectively (Pereira and Lupas, 2021). They are also widespread with respect to their phylogenetic distribution, occurring in all kingdoms of life, with a higher incidence however in individual branches, such as Actinobacteria, Cyanobacteria and Proteobacteria among the bacteria, Ascomycota and Basidiomycota among the eukaryotes, and Euryarchaeota among the archaea.

The cutoff for our analysis at 60% median internal sequence symmetry was chosen because only one natural β-propeller exceeding this value had been reported at the time and this protein was the one that had prompted this study in the first place. In fact, though, the range of internal sequence symmetries in β-propellers covers a continuum from less than 20% up to 100%. The focus on those with high internal sequence symmetry was therefore motivated by our effort to substantiate the independent amplification of a large number of diverse β-propellers. In the process, we also encountered a fair number of β-propellers that appear to be disrupted, indicating that the cycle may include an element of decay as well.

This prompts the question of the evolutionary mechanisms that govern this cycle.

β-Propellers are a closed fold, in which a variable number of repeats bring about an overall globular shape. The expectation from other repetitive globular folds is that an ancestral amplification at the origin of the fold was followed by an extended period of differentiation through descent with modification. This does not, however, appear to apply to all β-propellers (Chaudhuri, Söding and Lupas, 2008). In ancient β-propellers, like the one forming the G protein β subunit, blade similarity is indeed higher between equivalent blades in different homologs than within the same β-propeller, indicating the expected ancestral amplification, followed by gradual differentiation. In highly repetitive β-propellers, however, blade similarity is higher between blades of the same β-propeller than to any other blade, indicating a recent origin, independent of other β-propellers in the database.

We are thus faced with the question of the temporal balance between amplification and differentiation events. As we show for the WRAP proteins, even extremely similar β-propellers may have been amplified independently, being only homologous at the level of the blade, but analogous in the fully amplified form. Judging from the range of internal sequence symmetry in β-propellers, we are thus led to conclude that amplification events happen continuously in many lineages and are still ongoing today.

In our dataset, we encountered mostly forms that have been amplified globally from one single blade (92% of the dataset), although there was one instance of a form that had arisen from the amplification of a 2-bladed fragment, the first instance ever observed to our knowledge. The remaining 8% of the dataset, however, were instances of local amplifications within otherwise fully differentiated β-propellers. The most surprising of these resulted from the recent duplication of an exon in an ancient protein of bilateria, which appears to be species-specific even within the silk moths, where it occurred. We conclude that while global amplification is the dominant mechanism for new β-propellers, local amplification of individual blades is substantial and opens new evolutionary possibilities for otherwise fully specialised forms.

In mature β-propellers, particular lineages tend to prefer a well-defined number of blades (Kopec and Lupas, 2013). Since there is no counting system associated with the amplification process, we would anticipate that our set of recently amplified β-propellers would contain forms with more or fewer blades than preferred by that particular lineage. Indeed, we observe that repetitive β-propellers of a given lineage mostly have the blade number preferred by that lineage, but additionally manifest many larger and smaller ones. As seen from protein engineering studies, β-propellers with smaller numbers may reach a folded structure by oligomerisation (Smock *et al*., 2016; Afanasieva *et al*., 2019; Vrancken *et al*., 2020). These studies also show that β-propellers with larger numbers can fold to the preferred number by leaving one or more blades unstructured. While the presence of unstructured material is likely to be disfavoured and rapidly removed by in-frame deletion, it is less clear why oligomeric forms should be disfavoured, but the observation is that only a minute fraction of natural β-propellers are oligomeric. In either case, the newly arisen β-propeller will be likely to converge rapidly onto the preferred number of blades, either through further amplification or deletion.

In our dataset, there is a non-trivial number of β-propellers disrupted by frameshift and in-frame stop codons. Since these are highly repetitive forms, it would seem that they are decaying soon after their origin and might therefore represent proteins that were unable to converge on a form useful to the cell in a relevant timeframe. There is however also the possibility that their usefulness was present, but of limited duration, for example if they were involved in innate immunity or self-recognition. In this context, we note that there is a large number of disrupted β-propellers among the WRAP branch of WD40, which has been implicated in bacterial innate immunity, and among VCBS proteins, which may be involved in bacterial self-recognition (Dunin-Horkawicz, Kopec and Lupas, 2014).

Overall, this study has revealed a number of features that are not widely observed in protein evolution. Most conspicuously, the continuous amplification of new forms from smaller, subdomain-sized fragments is observed in fibrous folds, such as coiled coils (Hernandez Alvarez, Bassler and Lupas, 2019), and solenoids, such as TPR (Zhu *et al*., 2016), but not in globular folds. β-propellers appear to be unique in forming closed structures, yet undergoing continuous amplification from single blades. As a flip side of the continuous amplification, β-propellers also offer many examples for the decay of young protein coding genes, which either have never reached usefulness or have lost it within brief evolutionary time. For those that have not started decaying soon after their genesis, we are also afforded the opportunity to observe the accumulation of synonymous and non-synonymous mutations. In conclusion, β-propellers are an ideal model system in which to study the evolutionary cycle of proteins at warp speed.

## Acknowledgements

We would like to thank the Bioinformatics group of the Department of Protein Evolution at the Max Planck Institute for Developmental Biology, especially Dr. Laura Weidmann-Krebs and Hadeer Elhabashy, for stimulating discussions. JP would also like to thank the Schwede group at the Biozentrum of the University of Basel, especially Stefan Bienert and Gabriel Studer, for support on protein structure modelling with AlphaFold2 using af2@scicore. This work was supported by the Volkswagenstiftung [grant number 94810] and institutional funds of the Max Planck Society.

## Code and data availability

The code used and data generated for this project are available through: https://github.com/JoanaMPereira/RecentlyAmplifiedProps.

## SUPPLEMETARY

**Figure S1.**
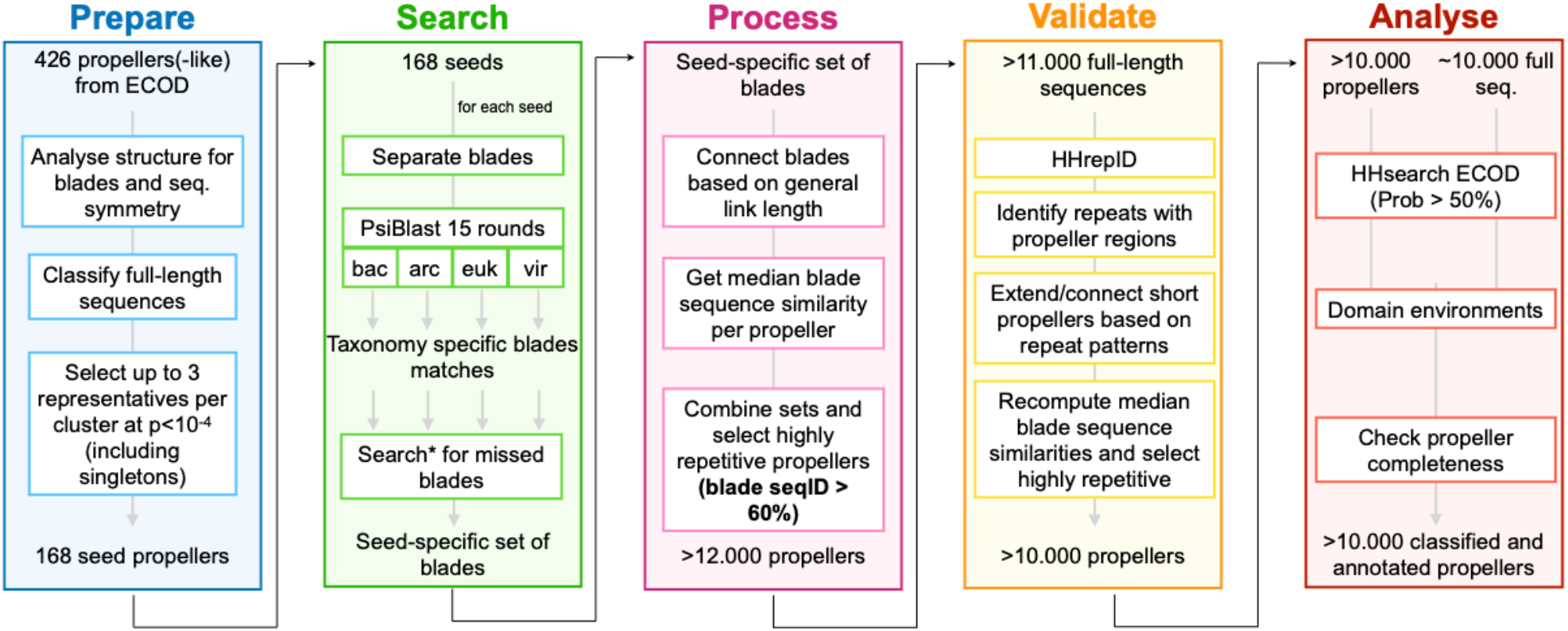
General workflow for the identification and annotation of protein sequences containing highly symmetric β-propellers. A detailed description of the procedure is given in the “Material and Methods” section.

**Figure S2.**
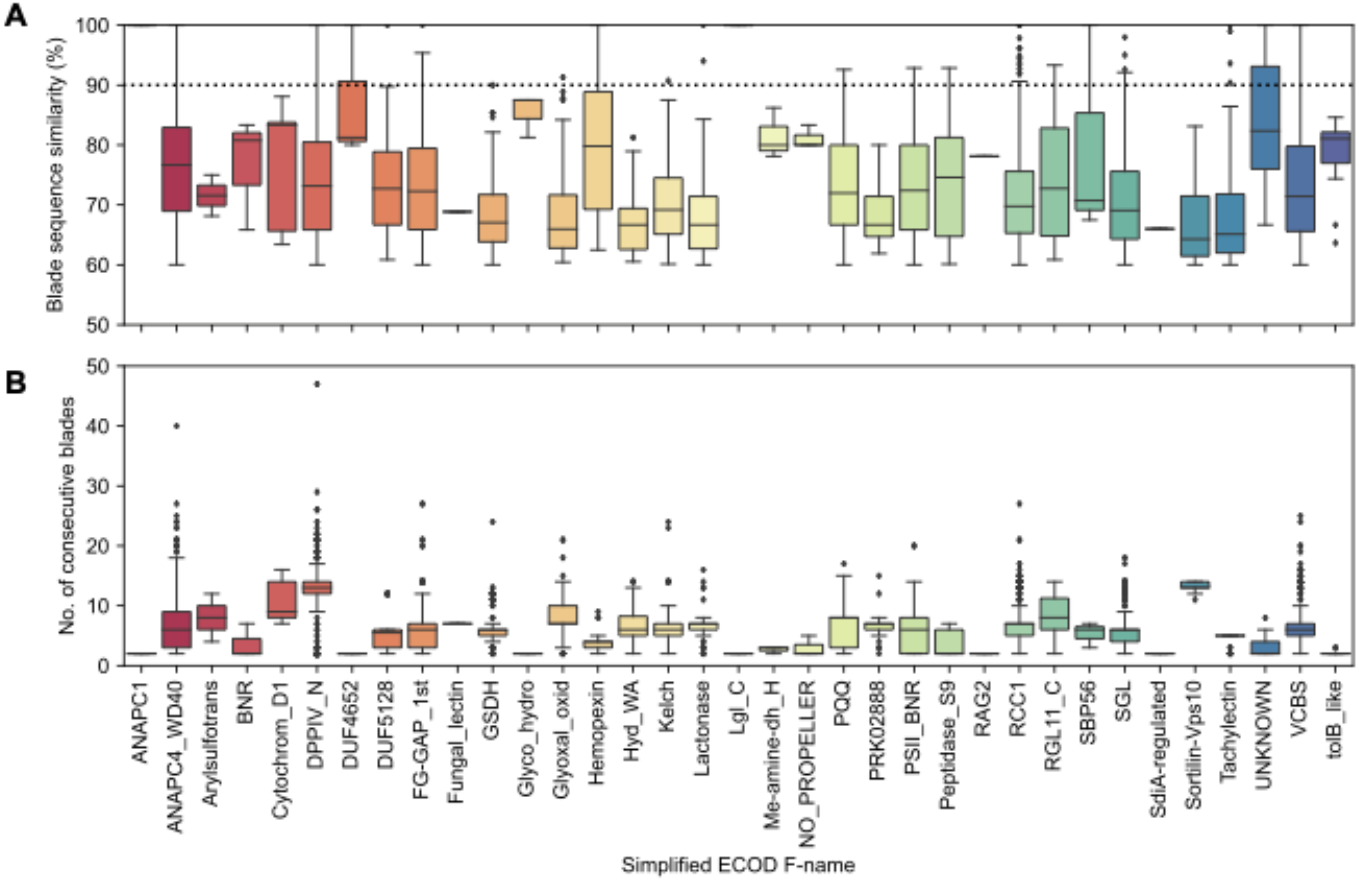
Boxplot of (a) the median sequence identity, and (b) the absolute number of consecutive highly similar blades for different β-propeller families.

**Figure S3.**
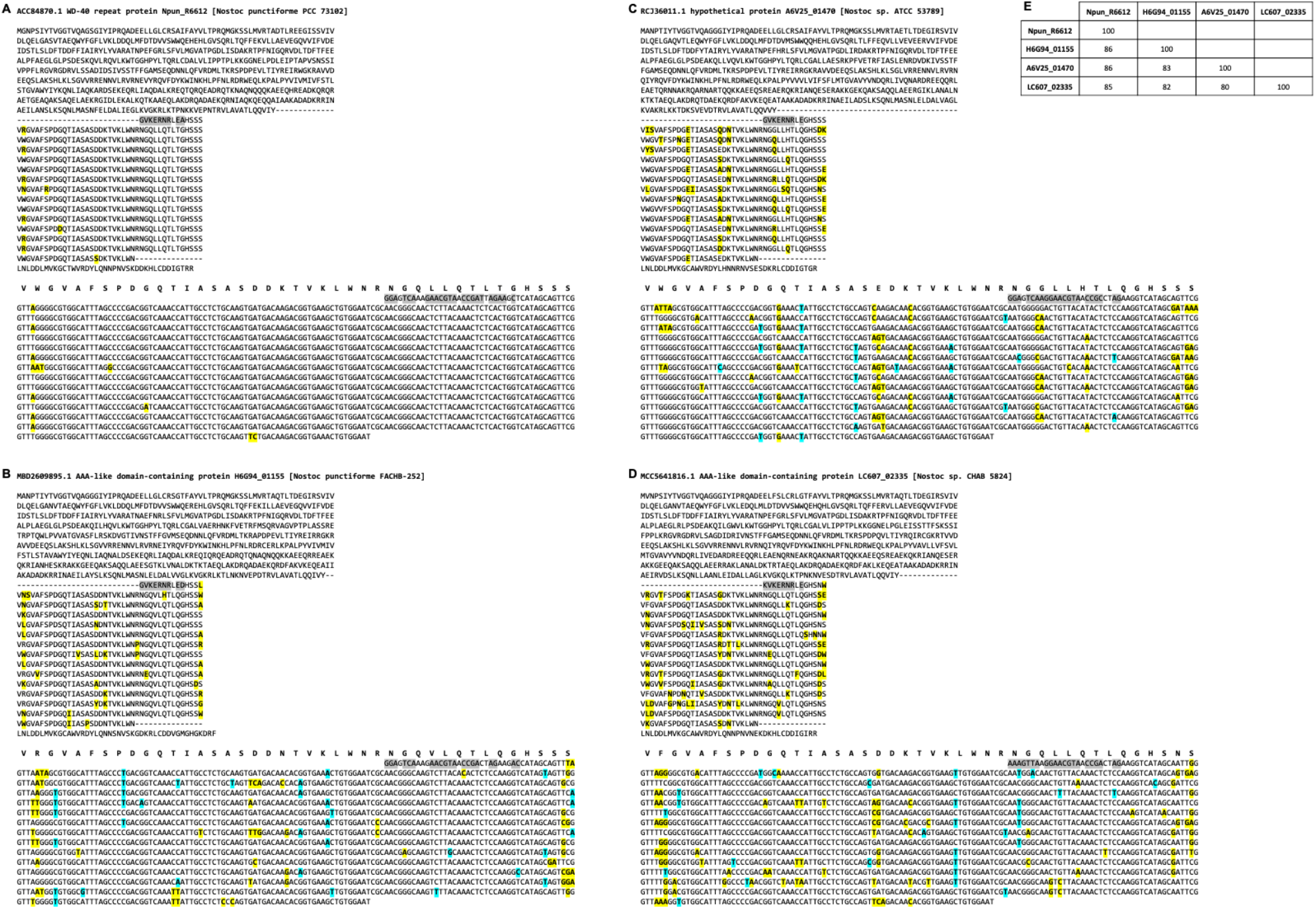
The four WRAP domain homologs in figure 9 and their pairwise sequence identities. (a-d) For each sequence, we show the alignment of the 14 blades with mutations highlighted in yellow and the corresponding nucleotide sequence with highlights as described in figure 2. The majority rule consensus of the β-propeller blades is shown above the nucleotide sequences. (e) Table of pairwise sequence identities.

**Figure S4.**
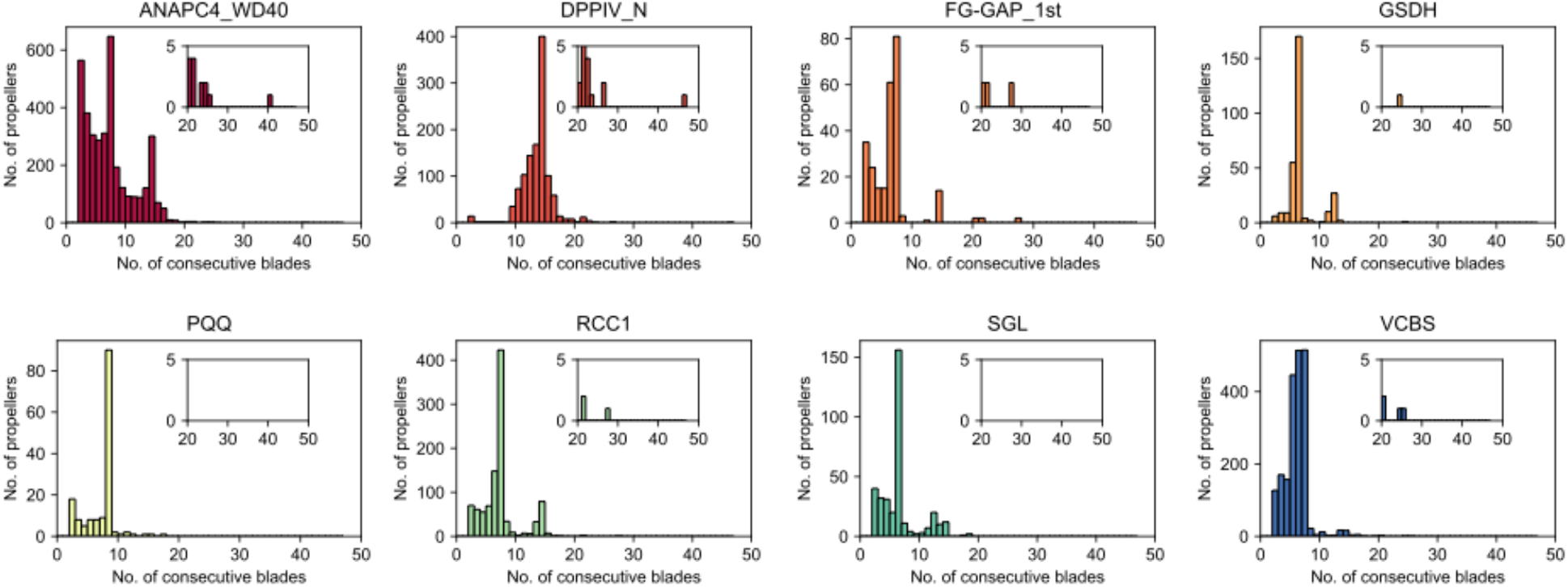
Histograms of the absolute number of consecutive highly identical blades for the globally highly repetitive β-propeller from the 8 most frequent families in the dataset.

**Figure S5.**
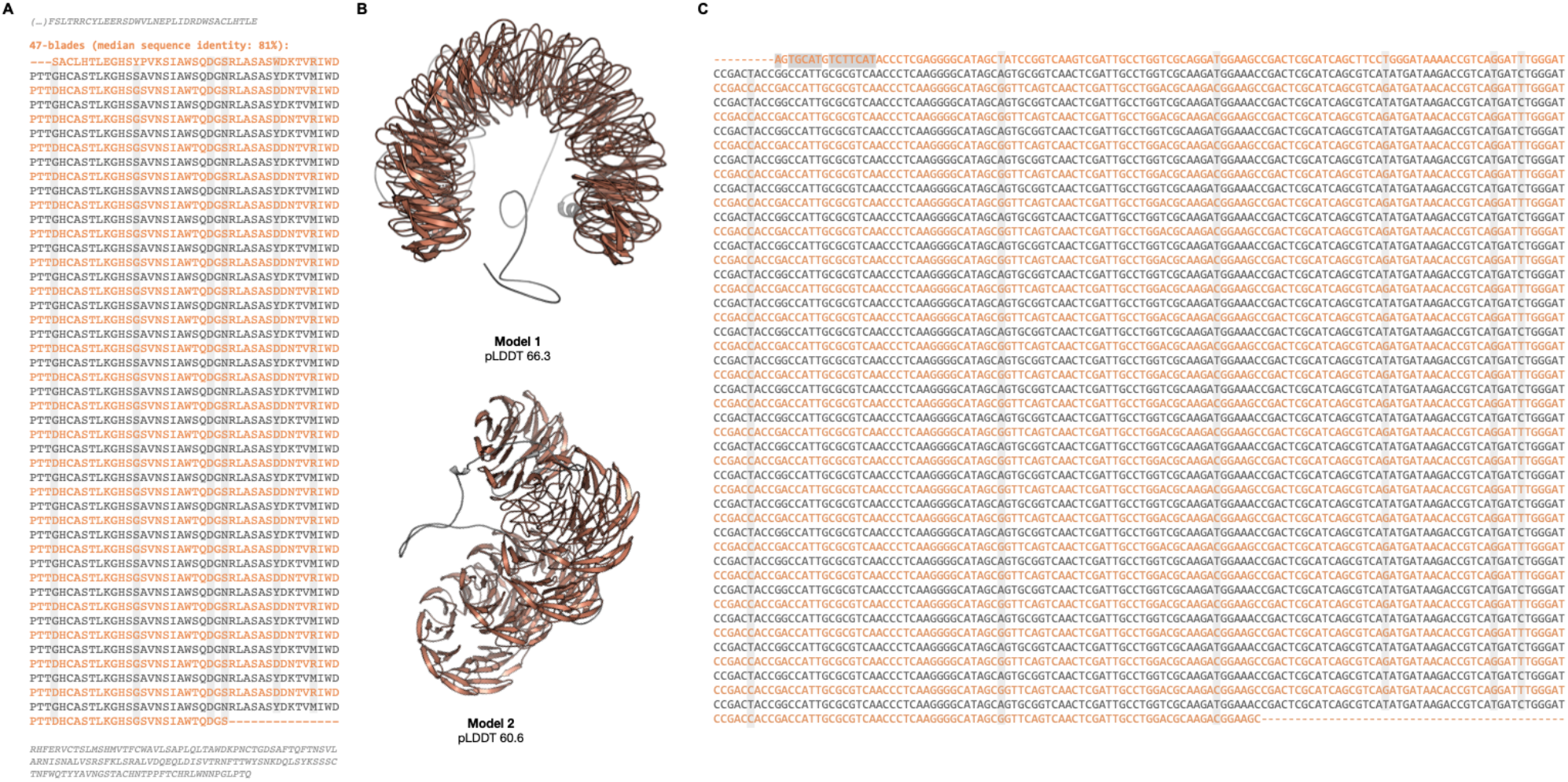
The 47-bladed globally highly repetitive β-propeller from *Penicillium camemberti* hypothetical NACHT nucleoside triphosphatase (CRL31138.1). (a) Annotated β-propeller sequence, highlighting the 2-step repeat pattern, plus the flanking non-propeller sequences. (b) The two best three-dimensional models of the β-propeller and immediate flanking regions predicted with AlphaFold v2.1.2 (Jumper *et al*., 2021). (s) The nucleotide sequence coding for the highly repetitive β-propeller region highlighting the same 2-step, intercalated, repeat pattern. In (a) and (b), point mutations are highlighted in grey.

**Figure S6.**
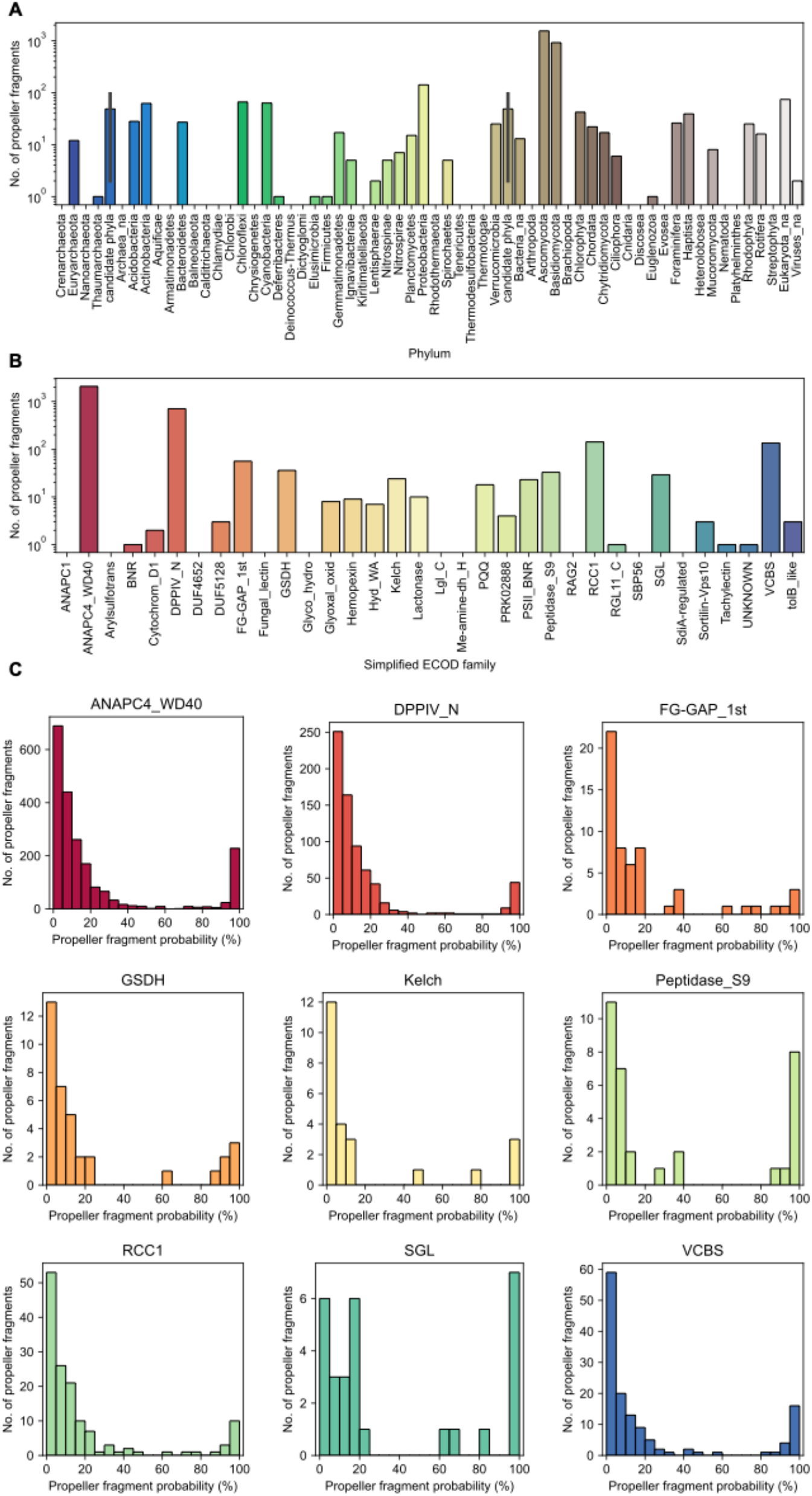
The overall distribution of putative β-propeller fragments. The number of fragments identified (a) per phyla and (b) highly repetitive β-propeller ECOD family. (c) Histograms of β-propeller fragments confidence (HHsearch probability) for the top nine families with the higher number of fragments.

**Figure S7.**
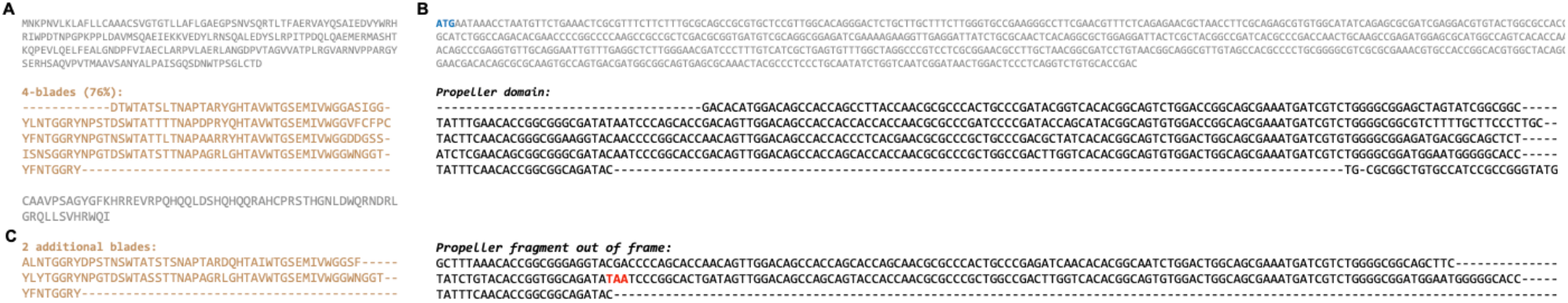
The β-propeller fragment in the neighbourhood of the globally repetitive β-propeller in hypothetical protein DMF28_08825 of an unknown Verrucomicrobia bacterium (PYL67592.1). (a) The protein sequence as deposited, highlighting the 4 highly repetitive blades. (b) The corresponding nucleotide sequence plus the 3’ genomic context. The start codon is highlighted in blue, and the in-frame stop codon in red. (c) The two additional blades hidden by a frame shift in the ORF that introduces the stop codon prematurely.

## Notes

https://github.com/JoanaMPereira/RecentlyAmplifiedProps

### Competing Interest Statement

The authors have declared no competing interest.

